# Conservation of differentiation but transformation of initiation in hippocampal neurogenesis

**DOI:** 10.1101/204701

**Authors:** Hannah Hochgerner, Amit Zeisel, Peter Lönnerberg, Sten Linnarsson

## Abstract

The dentate gyrus in the hippocampal formation is one of few regions in the brain where neurogenesis persists in the adult, and is therefore studied in the context of neurodevelopment and regenerative medicine. However, the relationship between developmental and adult neurogenesis has not been studied in detail. Here, we used extensive and unbiased single-cell RNA-seq to reveal the molecular dynamics and diversity of cell types in perinatal, juvenile and adult mice. We found clearly distinct quiescent and proliferating progenitor cell types, linked by transient intermediate states to neuroblast stages and fully mature granule cells. The molecular identity of quiescent and proliferating radial glia shifted after postnatal day 5, and was then maintained through postnatal and adult stages. A similar shift was observed for granule cells at P20. In contrast, intermediate progenitor cells, neuroblasts and immature granule cells were nearly indistinguishable at all ages. These findings demonstrate the fundamental continunity of postnatal and adult neurogenesis in the hippocampus, and pinpoint the early postnatal transformation of radial glia from embryonic progenitors to adult quiescent stem cells.

## Introduction

The dentate gyrus, part of the hippocampus, is involved in learning, episodic memory formation and spatial coding. It receives unidirectional input from the entorhinal cortex through the perforant path and projects to the CA3, forming the first step of the hippocampal trisynaptic circuit. Anatomically, the dentate gyrus is comprised of three layers: the cell-sparse molecular layer and the hilus, separated by the dense granule layer. The granule layer creates the characteristic V-shape in coronal and most sagittal hippocampus sections. It is densely packed with uniform, small granule cells whose dendrites extend into the overlying molecular layer, receiving input from the entorhinal cortex. Enclosed in the wedge-shaped granule layer and along the edge of the CA3 pyramidal layer is the hilus (polymorphic layer), where granule cell axons, called mossy fibers, project to the CA3. The excitatory granule cells are the most abundant cell type. In the molecular layer, a small number of interneurons are present, and the hilus contains a second type of principal excitatory neurons resembling CA3 pyramidal cells, called mossy cells. Further, a variety of GABAergic interneurons have been described, such as the pyramidal basket cell, enriched at the interface of the granule layer to the hilus, known as the subgranular zone, which provide inhibitory control of granule cell activity (schematic Fig. 1e, reviewed in (Amaral et al. 2007)).

The dentate gyrus develops relatively late, and several key developmental events occur postnatally. The majority of granule cells are born perinatally, with significant numbers of cells generated until around postnatal day ten (P10). This takes place in two distinct waves of neural progenitor cell migrations (Altman & Bayer 1990; Piatti et al. 2006). In rat, the first migration originates from the primary dentate neuroepithelium, and generates the outer granule cell layer by E19. Early postnatally, a second migration forms the tertiary dentate matrix, with radial migration of progenitors contributing to the inner granule cell layer (Fig 1A). Radial glia cells (also called radial astrocytes) are neuronal stem cells during embryonic development (Malatesta et al. 2000; Malatesta & Götz 2013), and are still abundant in the first postnatal weeks. Radial glia-like cells differentiate into proliferating progenitors (neuronal intermediate progenitor cells or nIPCs) (Kriegstein & Alvarez-Buylla 2009) and exit the cell cycle to become migrating neuroblasts, then settle in the granule cell layer and mature around postnatal week 3 into fully functional excitatory granule cells.

**Figure 1.**
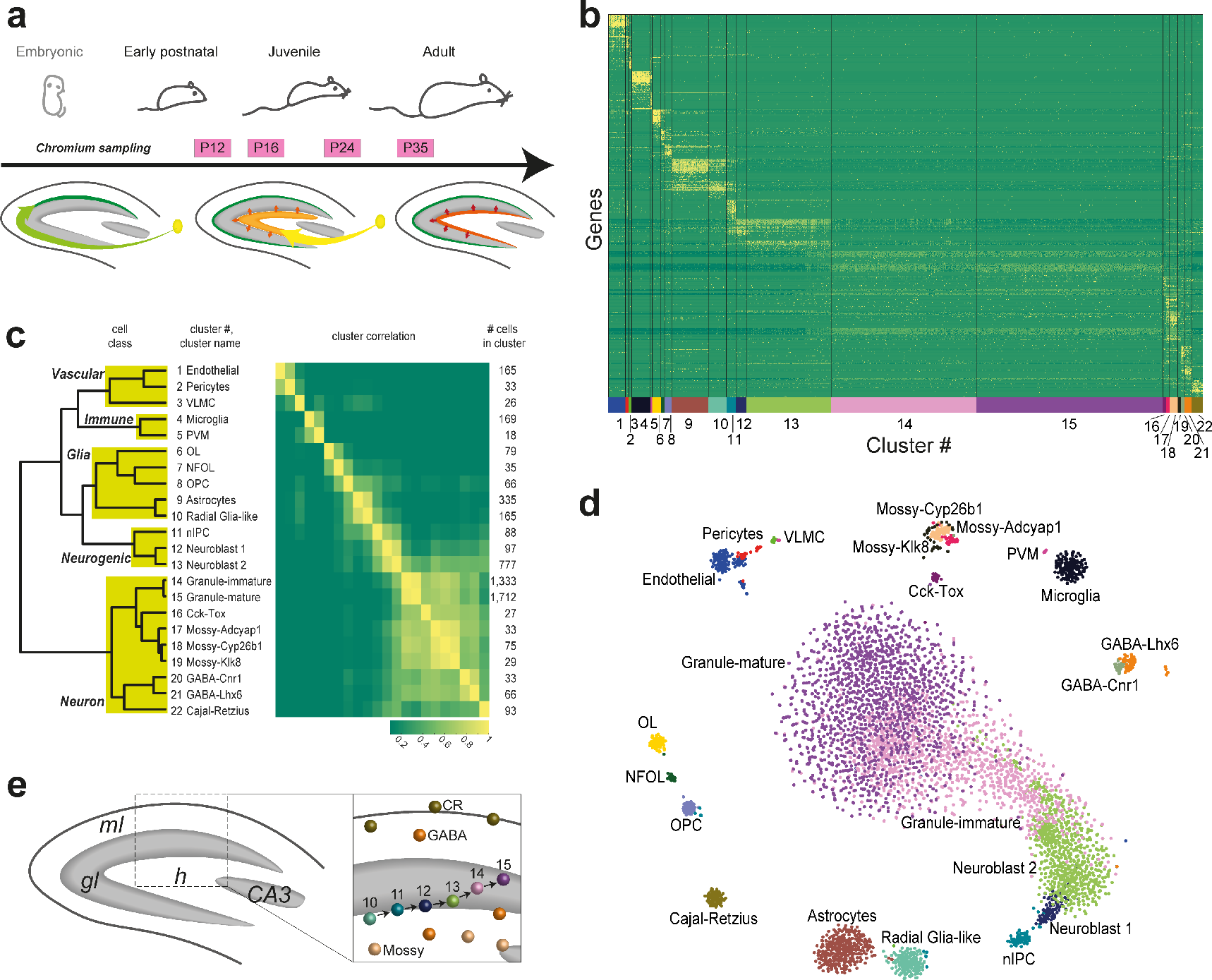
Transcriptional architecture of the mouse dentate gyrus. (a) Sampling time points along postnatal development. The dentate gyms schematics (adapted from (Piatti et al. 2006)) depict waves of granule neurogenesis. Left: Perinatal migration of neural progenitor cells (NPCs) from the primary dentate neuroepithelium (yellow), generating outer granule layer (green). Middle: Early postnatal migration to form the tertiary matrix (orange), with NPC radial migration into the inner granule layer. Right: Adult neurogenesis from NPCs retained in the subgranular zone (red). (b) Heatmap of 22 clusters from 5454 cells (all sampled time points), represented by their top marker genes (648 genes), expression normalized by gene. (c) Dendrogram and correlation matrix of the 22 clusters identified. Boxes underlaying the dendrogram indicate main cell classes. (d) t-SNE visualization of 5454 cells, colored by cluster annotation. (e) Structure of the dentate gyrus and previously described spatial organization of cell types in relation to clusters identified in this dataset (*ml* – molecular layer, *gl* – granule layer, *h* – hilus, *CA3* – hippocampus CA3 pyramidal layer).

In adult mammals, radial glia-like cells are neuronal stem cells in the subventricular zone (Doetsch et al. 1999) and continue to generate granule cells from the dentate gyrus subgranular zone (Seri et al. 2001; Steiner et al. 2004; Garcia et al. 2004). Thus, a small number of progenitor cells in the subgranular zone continue to generate new granule cells throughout life, albeit at low rates (Altman & Das 1965; Kaplan & Hinds 1977; Eriksson et al. 1998; Gould et al. 1999; Gage 2000; Kempermann et al. 2004). The process is believed to contribute to learning and memory formation, and is sensitive to stress, antidepressants as well as enrichment and exercise (Kempermann et al. 1997; Kempermann et al. 1998; Gage et al. 1999; Gonçalves et al. 2016). Adult-generated granule cells undergo a process of maturation, lasting several weeks, that is strongly dependent on input from the entorhinal cortex. In fact, the process of neuronal integration bears a clear resemblance to maturation of granule cells during early postnatal development (Espósito et al. 2005; Laplagne et al. 2006; Overstreet-Wadiche et al. 2006). Both early postnatal and adult neurogenesis in the dentate gyrus are highly controlled processes, where cells go through clearly distinct stages of differentiation and maturation. Morphologically, the adult neuronal stem cell (NSC) closely resembles the postnatal radial glia-like cell, with a single radial process extending through the entire granule cell layer (Götz et al. 2016). In the adult, NSCs are label-retaining and persist in a quiescent state, occasionally giving rise to a proliferating progenitor. Upon differentiation in the adult, newborn neuroblasts go through a several weeks-long process of maturation, acquiring mature membrane properties after about three to four weeks (Overstreet-Wadiche & Westbrook 2006). While adult NSCs and early postnatal radial glia-like cells show many similarities, they act in different environments, perhaps requiring different regulatory mechanisms to maintain their neurogenic potential (Götz et al. 2016). However, it is not currently known whether NSCs are simply radial glia-like cells that persist, or whether there are any molecular properties that make adult NSCs unique. Similarly, it is not currently known if early postnatal and adult neurogenesis proceed through identical intermediate states, or if the adult process is different beyond timing (Overstreet-Wadiche et al. 2006) from the initial postnatal waves.

More fundamentally, both early postnatal and adult neurogenesis involve very closely related cell types, which have been difficult to separate based on molecular markers. For example, glial fibrillary acidic protein (GFAP) is often used to isolate NSCs, but is also strongly expressed in astrocytes. The question arises whether astrocytes, radial glia-like cells, NSCs and nIPCs are distinct cellular states with clearly delineated molecular properties (such as transcription factors).

Recent work using single-cell RNA-seq and targeting the neurogenic populations have instead suggested that neurogenesis forms a continuum of molecular states (Shin et al. 2015; Habib et al. 2016). However, we suspected that the limited number of cells in those studies, and the targeted sampling using specific markers, might have masked the presence of distinct cell types. We therefore performed larger-scale single-cell RNA-seq and unbiased sampling to delineate the cellular states along the granule cell lineage. We examined perinatal and juvenile development as well as adult mice, and used two complementary platforms to rule out batch effects and obtain both wide and deep sampling of cells and time points. We unraveled a branching lineage including all major neural cell types: astrocytes, oligodendrocytes, granule cells and hippocampal pyramidal neurons or mossy cells. We observed a clear sequence of distinct steps that constitute granule cell development from quiescent stem/progenitor cells to mature neurons, and we validated these findings using *in situ* hybridization. Comparing cells across time from E16.5 to P132, we demonstrate a perinatal transformation of neurogenesis in the dentate gyrus, from an embryonic to a postnatal configuration. During the second postnatal week, radial glia shifted sharply from an embryonic to an adult state, and during the same time, cycling intermediate precursor cells shifted from a less to a more neurogenic state. A similar step-wise maturation of the granule cells was observed around the third postnatal week, establishing a distinct post-adolescent granule cell identity. In contrast, neuroblasts and immature granule cells were indistinguishable at all ages. Interestingly, the distinct fates of granules cells and hippocampal pyramidal neurons emerged from the shared postmitotic neuroblast state, implying that their fate was determined only after they exited the cell cycle.

## Results

### Cell census of the developing dentate gyrus

In order to address unanswered questions about the postnatal development of the dentate gyrus, we performed single-cell RNA-seq on microdissected mouse dentate gyrus. Dataset A comprised four time points, postnatal days 12 (P12), P16, P24 and P35, including a total of 5,454 cells using droplet-based single-cell RNA-seq (10X Genomics Chromium; see Methods). To rule out batch effects, we include a set of experiments (Dataset B) using an orthogonal single-cell RNA-seq technology (valve-based microfluidics chips; Fluidigm C1), sampled at 17 developmental time points, with 2,303 cells. Based on these initial findings, we acquired a third larger dataset (Dataset C, 10X Chromium), extending to earlier and later timepoints, comprising 24,185 cells from E16.5, P0, P5, P18, P19, P23, P120 and P132. We analyzed the datasets separately because they were obtained using different technologies (A and C vs. B) and kit versions (A vs. C). All datasets are summarized in Supplementary Figure 1. We provide an online browsable resource (at http://linnarssonlab.org/dentate) where gene expression can be visualized in individual cells and clusters of all three datasets.

Focusing first on Dataset A (Fig. 1a), graph based clustering analysis (Markov clustering (van Dongen & Abreu-Goodger 2012) of the mutual k-nearest neighbor graph; see Methods) revealed 22 distinct types of cells (Fig. 1b), which were arranged in four major categories (Fig. 1c): vascular (endothelial, pericytes, and vascular leptomeningeal cells), immune (microglia and perivascular macrophages), glial and neuronal cell types.

Visualizing these cells using t-Distributed Stochastic Neighbor Embedding (t-SNE (Maaten & Hinton 2008), Fig. 1d) remarkably revealed the entire architecture of the postnatal developing dentate gyrus in detail (Fig. 1e). At center, dominating the structure was the granule cell lineage starting with radial glia-like cells (RGLs), via neuronal intermediate progenitor cells (nIPCs), two neuroblast stages (Neuroblast 1 and 2), immature and mature granule cells. RGLs were closely related to, but clearly distinct from, astrocytes. Oligodendrocytes also formed a three-stage lineage (OPC, NFOL, OL(Marques et al. 2016)), while all remaining cell types formed distinct and separated clusters. Enriched in P10 and P16 samples we found Reelin (*Reln*)-expressing Cajal-Retzius cells, which are required for proper morphogenesis and migration in both the cortex and the hippocampus. We found that in addition to the more broadly expressed *Reln*, Cajal-Retzius cells highly specifically expressed homeobox domain genes such as *Lhx1* and *Lhx5* (Suppl. Fig. S2)(Hawrylycz et al. 2012), as well as cell death regulator genes *Trp73* (encoding p73) and *Diablo*.

### Radial Glia-like Cells are different from astrocytes and neuronal Intermediate Progenitor Cells

We first sought to identify the key cell types of early neurogenesis, that is RGLs, nIPCs, neuroblasts and granule cells (Fig. 2). RGLs were identified by their close similarity to astrocytes (*Gfap*, *Hes5* and *Sox9*), and distinct expression of recently described neuronal stem cell marker *Lpar1* (Walker et al. 2016) (also expressed in oligodendrocytes, Fig 3c). They almost completely lacked expression of the aquaporin *Aqp4*, characteristic of mature astrocytes. They expressed both *Nes* (nestin) and *Prom1* (also known as CD133), commonly used stem cell markers, although both genes were more highly expressed in endothelial cells (Fig 3c). The complete absence of cell cycle genes such as *Cdk1*, *Top2a* and *Aurkb* showed that these cells are quiescent, in contrast to the actively dividing nIPCs.

nIPCs were an actively cycling progenitor population, with a strong cell cycle machinery signature (e.g. *Cdk1*, *Aurkb*, *Top2a* and many more). In addition, these cells had taken on an early neuronal fate, expressing the known neurogenic transcription factors *Neurog2*, *Eomes* and *Neurod4*. This is consistent with previous findings in the developing neocortex, where *Eomes* (also known as *Tbr2*) is not expressed by RGL cells; only when entering a proliferating intermediate progenitor state, *Tbr2* is switched on (Noctor et al. 2008). In the t-SNE plot, which arranges cells based on their overall transcriptional similarity, nIPCs were located at the root of a trajectory leading to mature granule cells (Fig 1d).

We further found two neuroblast populations, here defined as the first step of differentiation after nIPCs exit the cell cycle. The first population, Neuroblast 1, retained *Eomes* expression (shared with nIPCs), *Tac2* and *Calb2* (which encodes Calretinin), while the second, Neuroblast 2, expressed *Gal* and both populations shared expression of *Sox11* and doublecortin (Dcx). Both neuroblast stages also expressed *Igfbpl1*, a marker that we found to be more specific compared to the widely used *Dcx* and *Sox11* markers (Fig. 2). Together, our findings identify these cells as two sequential steps in the maturation of migrating neuroblasts (Hsieh 2012).

In order to identify possible transitions between these cell types, we reanalyzed astrocytes and the early neurogenesis clusters (RGL, nIPC, Neuroblast 1 and 2) using a graph representation that shows pairs of mutual nearest neighboring cells in gene expression space (mutual KNN graph, Fig. 2a). In this representation, cells are linked by graph edges when they are each other’s nearest neighbors in the high-dimensional gene expression space, suggesting possible transitions in expression space. Conversely, the absence of edges suggests disallowed paths through expression space. Graph edges confirmed the order of the trajectory from RGLs, via nIPCs to Neuroblast 1 and then Neuroblast 2. Assuming parsimony in gene expression, it also confirmed that neurogenesis proceeds essentially linearly through a sequence of steps, and that the two neuroblast populations are sequential, not parallel. It also demonstrated the close relationship; yet clear distinction, between astrocytes and RGLs.

With dentate gyrus neurogenesis dissected into several molecularly distinct steps, we performed an unbiased search for cell type-specific gene expression. Focusing on transcription factors, reassuringly we rediscovered previously known key regulators *Hes5* (radial glia-like cells and astrocytes), *Neurog2* (nIPCs), *Neurod4* and *Eomes* (Neuroblast 1), and *Sox11* (Neuroblast 2). In addition, using the wide scope of the current study, we identified several highly specific markers and present an expression sequence of single and combinatorial genes (Fig 2a-c, 3c, Suppl. Fig. S3, Suppl. Table 1, 2), marking the progression from RGL, via nIPC to Neuroblast 1 and 2. For instance, we found the transcription factor *Tfap2c* specifically expressed in quiescent radial glia-like cells and nIPCs and propose it as a marker in the context of early neurogenesis. *Vnn1*, *Rhcg* and *Wnt8b* are further examples of genes more narrowly expressed in RGL cells, than commonly used markers.

**Figure 2.**
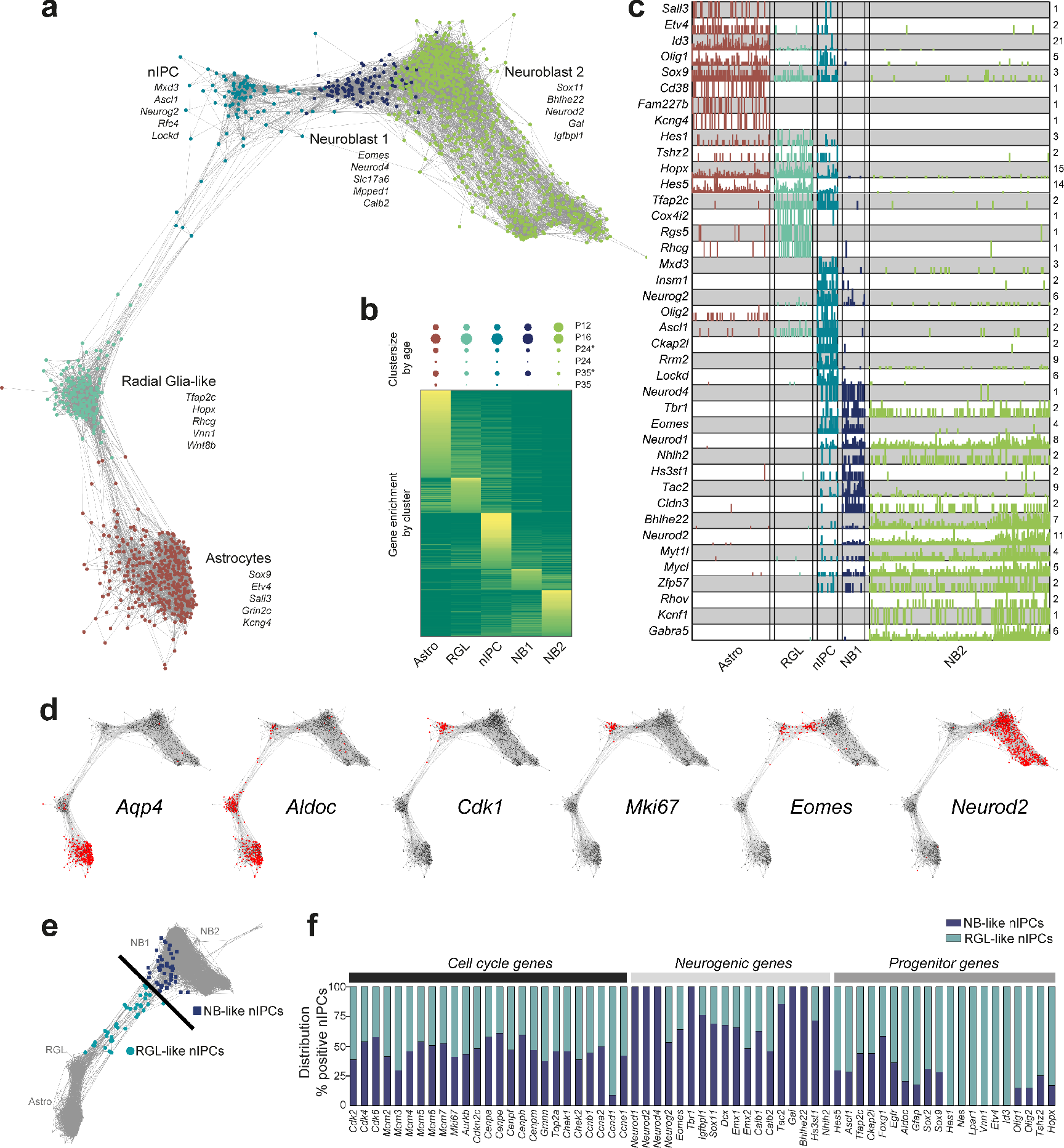
Dissecting the molecular progression through early neurogenesis. (a) Mutual KNN graph of astrocytes and early neurogenesis-related clusters (radial glia-like (RGL), neuronal intermediate progenitor cells (nIPC), Neuroblast (NB) 1 and 2)). Nodes represent cells, edges connect mutual *k*-nearest neighbors (*k* = 40). Examples of markers enriched in each type are indicated. (b) Heatmap of genes with differential expression during early neurogenesis, sorted by their peak cluster. Full list of genes and expression provided as Supplementary Table 1. Above, the frequency of each cluster per age is depicted by circles (proportional to area). P24* and P35* include an estimated 85% of P24 or P35 cells, while 15% are contributed from the P16 and P12 samples, respectively, due to sex-based sample separation inaccuracy, see Methods. (c) Single-cell expression of representative top enriched markers for each stage (number of molecules). Numbers on the right indicate maximum detected expression. (d) Mutual KNN graphs as in (a), stained by the expression of genes indicated. (e) Mutual KNN graph with nIPC-specific genes removed (Suppl. Fig. S3). nIPCs were divided by their position along the transition between RGL and NB, as indicated by the line. (f) Distribution of the percentage of NB-like or RGL-like nIPCs positive for genes associated with cell cycle, neurogenesis or progenitor cells.

A fundamental outstanding question in dentate gyrus neurogenesis concerns the distinction between quiescent and active progenitors/stem cells. Is quiescence a distinct transcriptional state, or is it just the absence of cell division? What is the nature of the transition from quiescence to active division, and how is it related to neuronal commitment? As mentioned, nIPC was the only cluster of cells that was proliferating, but it also already expressed neurogenic transcription factors. Both nIPCs and neuroblasts were depleted with age (Fig 2b), whereas RGLs were maintained. But how could RGLs be maintained in the absence of cell division? We speculated that a small population of proliferating RGLs might have been clustered with nIPCs, which would not be unexpected since cell division involves the activation of large numbers of genes, which would dominate and drive clustering. To further dissect the proliferating nIPCs, we removed all genes expressed specifically in nIPCs (which would include cell cycle genes), but not those shared with either RGLs or neuroblasts. We then asked if individual nIPC cells now were more similar to neuroblasts or to RGLs. After recomputing the KNN graph, we found that most nIPCs were now intermingled among the earliest neuroblasts (Fig 2e, Suppl. Fig. S4), consistent with their commitment to neuronal differentiation. However, interestingly, a small number of cells remained close to RGLs in gene expression (expressing e.g. *Hes5*, *Ascl1*, *Lpar1* and *Nes*), did not express most neurogenic transcription factors (*Neurod1*, *Tbr1*, *Eomes*, *Neurod2*, *Neurod4*) but were still actively proliferating (many genes, including *Cdk1*, *Top2a*, *Cenpe*, *Aurkb*, *Mcm2-8* and *Mki67*). We confirmed this observation by splitting nIPCs into RGL-like and neuroblast-like subsets (Suppl. Fig. S4) and measuring the enrichment of progenitor-, cell cycle-, and neuroblast-specific genes (Fig. 2e-f). This analysis demonstrates the existence of a small population of dividing radial glia-like cells that have not entered the neurogenic program. Thus RGLs were mostly quiescent, but could be maintained by occasional proliferation without differentiation (or, alternatively, by occasional asymmetric cell division). This is also consistent with the observation in the subventricular zone that most stem cells are quiescent (91.4% based on the hGFAP:GFP reporter (Ponti et al. 2013)).

To validate genes expressed specifically in each of the steps of early neurogenesis (Fig. 2) and locate cell types in the tissue, we performed multiplexed RNA staining of the dentate gyrus (Fig. 3a). *Ascl1*+/*Tfap2c*+/*Cdk1*− RGLs were almost entirely confined to the subgranular zone at all ages, as were the proliferating *Ascl1*+/*Tfap2c*+/*Cdk1*+ nIPCs (Fig. 3b, top row). Neuroblasts (*Igfbpl1*+/*Eomes*+ (NB1) or *Eomes*- (NB2)) accumulated just above the SGZ, spanning almost the entire thickness of the granule cell layer at P10, but were transient and replaced by mature granule cells (*Plk5*+/*Eomes*-) at the later ages (Fig. 3b & d, second and third rows). These findings confirm the existence of distinct RGL, nIPC and neuroblast cell types, and reveal their spatial and temporal dynamics (Fig. 3d). Further, we showed that the separate or combinatorial expression of these genes labeled the cell types more specifically than previously used markers such as *Gfap*, *Hes5* or *Sox2* (shared with astrocytes), *Prom1* and *Nes* (shared with vascular endothelial cells) or *Lpar1* (expressed also in oligodendrocytes) (Fig. 3c).

**Figure 3.**
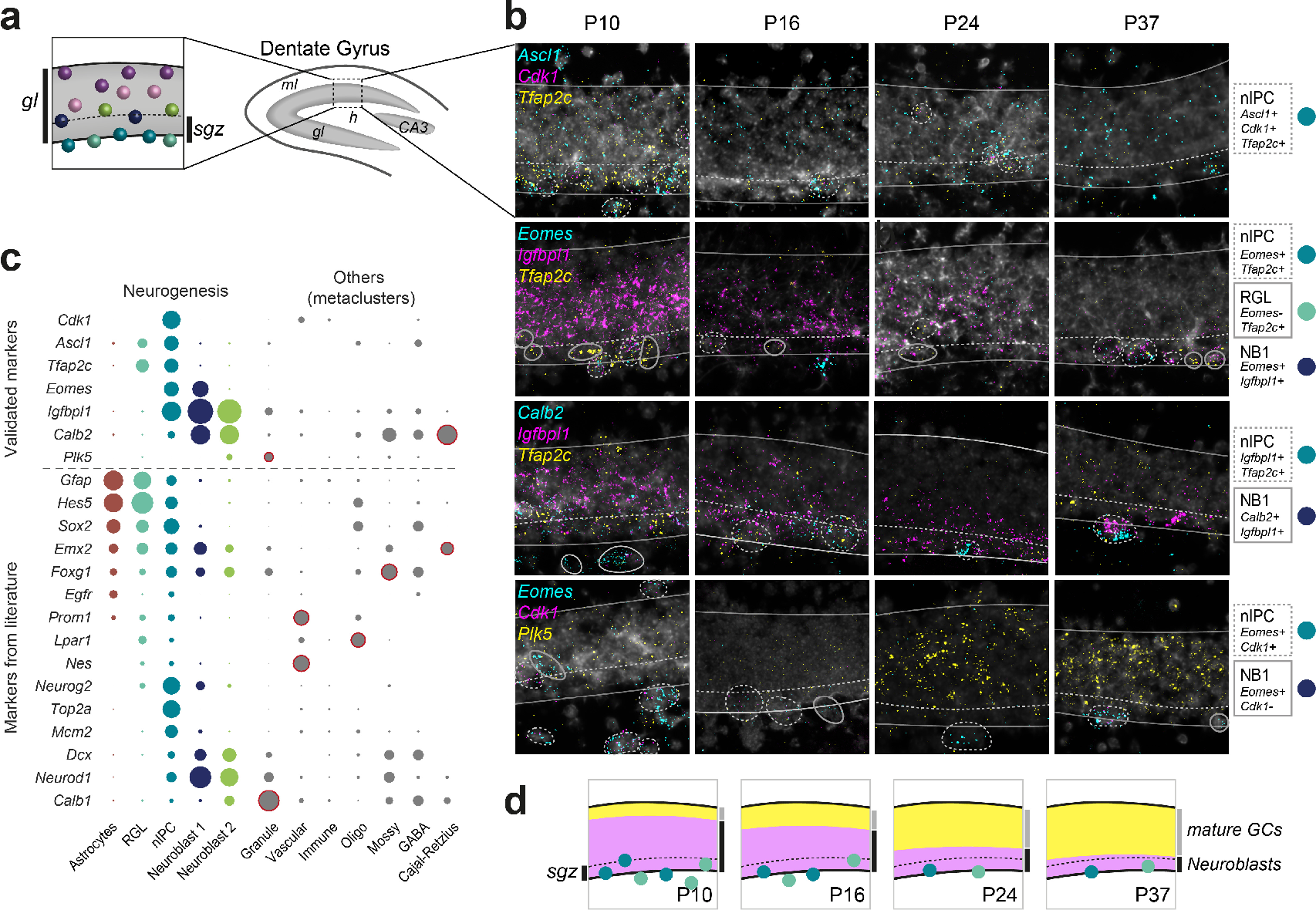
Validation and spatial localization of early neurogenic markers across age. (a) Schematic of the dentate gyrus, and zoom-in on the granule layer (*ml* – molecular layer, *gl* – granule layer, *h* – hilus, *sgz* – subgranular zone). (b) RNAScope multiplex *in situ* hybridizations of dentate gyrus sections from mice aged P10, P16, P24 and P37. Probes used against the marker genes are indicated in each row. Solid outlines depict the granule layer, including subgranular zone (below dashed line), as in (a). Dashed and solid circles highlight cells (approximate) of particular interest, as described on the right of the panels. (c) Proportion of cells positive for markers validated here (top), and other common markers from literature, for astrocytes and cell types in early neurogenesis (left) and other cell types summarized by class (right, grey circles). The circle areas are proportional to the frequency of cells with expression >0 within the cluster. ‘Other’ cell class expression circles are outlined in red, if the gene expression is enriched compared to neurogenic cell types. (d) Schematic summarizing neurogenesis validation in the granule layer across time points. RGL and nIPC are represented by colored circles as in (c), yellow shaded area illustrates area enriched in mature granule cells (*Plk5* staining in (b)) and pink shaded area enriched in neuroblasts (*Igfbpl1* staining in (b)). *sgz* – subgranular zone (below dashed line).

### Rapid maturation of granule cells around postnatal week 3

Noting the rapid maturation of neuroblasts into mature granule cells between P16 and P24 (compare *Igfbpl1* and *Plk5* stainings in Fig. 3b), we next focused on the later stages of neurogenesis. As granule cells matured, they transitioned from the still glia-like neuroblasts (Fig. 1d-e) into an immature granule cell state and finally to mature granule cells (Fig. 4a). This transition was characterized by a shift of gene expression most pronounced between the neuroblast to immature stage, but also from the immature to mature stage (Fig 4b, Suppl. Table 3, 4). Comparing developmental time points, we noted that mature granule cells appeared rather abruptly sometime between P16 and P24 (Fig 4c). This finding was confirmed in tissue by staining for *Fxyd7* (neuroblasts), *Prox1* (all stages), and *Ntng1* (mature granule cells) (Fig 4d-f). Although a small number of *Ntng1*-staining cells were observed prior to P16, their numbers greatly increased at P24 and filled most of the granule cell layer at P37.

**Figure 4.**
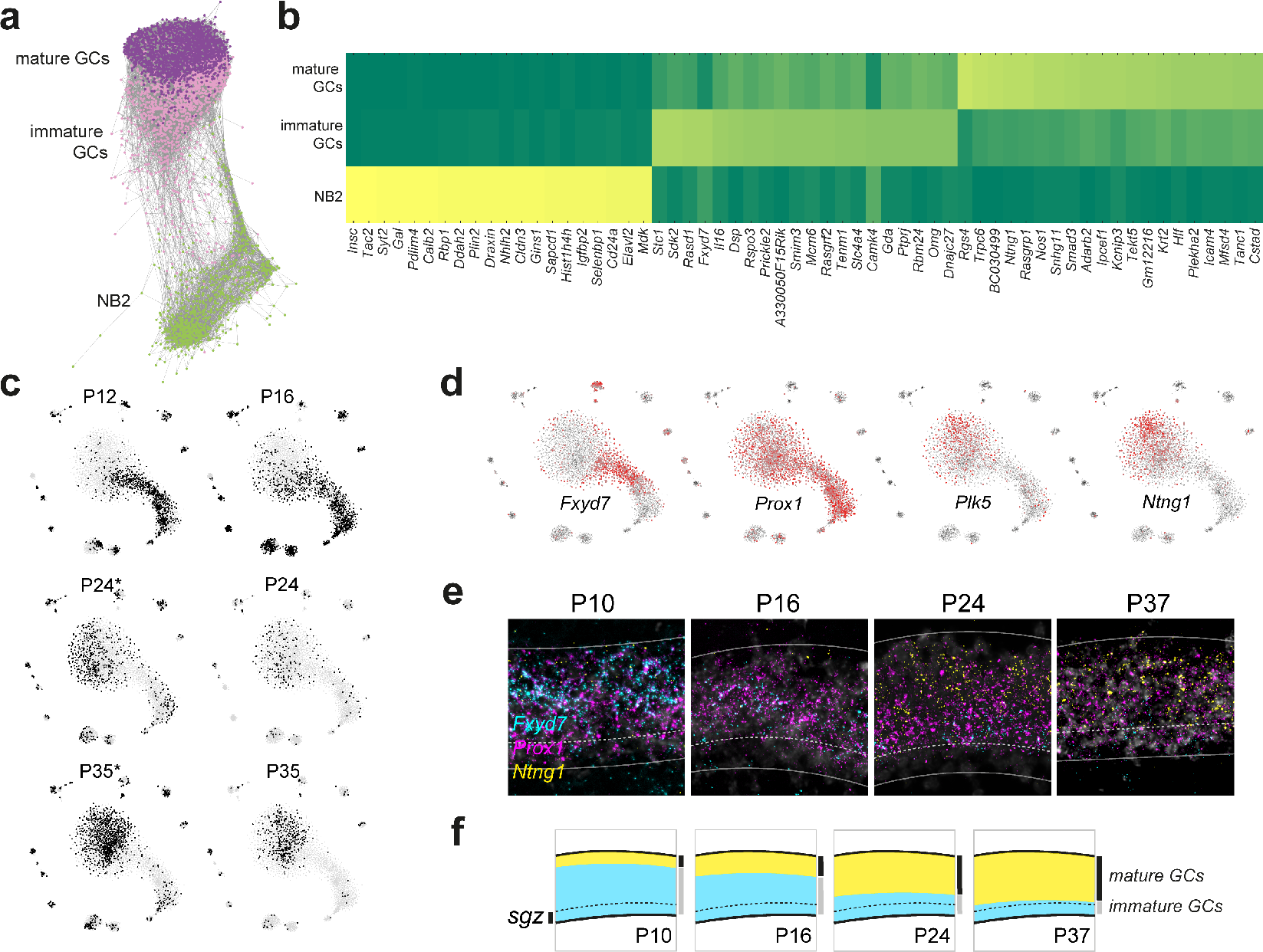
Granule cell maturation. (a) Mutual KNN graph (*k* = 40) of Neuroblast 2 (NB2), immature and mature granule cells (GCs). Nodes represent cells, edges connect mutual nearest neighbors. (b) Heatmap of genes with differential expression between the maturing cell types, sorted by their peak cluster. (c) t-SNE (as in Fig. 1d), stained by contribution from each sampling time point. P24* and P35* include an estimated 85% of P24 or P35 cells, while 15% are contributed from the P16 and P12 samples, respectively, due to sex-based sample separation inaccuracy, see Methods. (d) t-SNE (as in Fig. 1d), stained by expression of four markers; *Prox1* (pan-granule), *Fxyd7* (immature-enriched), *Plk5* and *Ntngl* (mature-enriched). (e) RNAScope multiplex *in situ* hybridizations of dentate gyrus granule layer (as in Fig. 3a-b), from mice aged P10, P16, P24 and P37. (f) Schematic summarizing maturation of granule cells, with immature granule cells (GCs) enriched in blue shaded area (*Fxyd7*+), mature granule cells enriched in yellow shaded area (*Ntng1*+, *Plk5*+ (Fig. 3b)). *sgz* – subgranular zone.

We had taken precautions to eliminate batch effects, including analyzing different ages in the same cell preparation and droplet emulsions. However, we were still concerned that our findings of an abrupt shift from immature to mature granule cells around postnatal week 3 might have been spurious. To address this concern, and to extend the previous findings to a higher temporal resolution, we performed a second set of experiments using an orthogonal single-cell RNA-seq technology (Dataset B, Fluidigm C1). We sampled 2303 single cells at 17 developmental time points and extending into adulthood, from P8 to P68 (Suppl. Fig. S1). We performed clustering and identified cell types independently in this separate dataset, which confirmed the findings previously obtained by droplet microfluidics (Suppl. Fig. S5). We confirmed the existence of distinct astrocyte, RGL, nIPC and neuroblast cell types, as well as the distinction between immature and mature granule cells. At P50 and P63, we enriched for neuronal stem cells by exploiting the hGFAP:GFP reporter mouse (Pastrana et al. 2009). We observed that *GFP*+ cells did not necessarily express endogenous *Gfap*, and included not only astrocytes and RGLs, but also PVMs, OPCs, OLs and neuroblasts (Suppl. Fig S5). This demonstrates not only that a broader expression in several cell types needs to be considered to avoid confounding results when using hGFAP:GFP to study adult neurogenesis, but also the power of single-cell RNA-seq to dissect and resolve the heterogeneity of complex samples.

Taking advantage of the denser temporal sampling, we not only confirmed the rapid maturation of granule cells around the third postnatal week, but also found that it occurs in a very short time interval (Suppl. Fig. S1). Immature granule cells virtually disappeared after P18, whereas mature granule cells appeared abruptly at P20 (which also agreed with our *in situ* validation, Fig. 4e). A similar abrupt disappearance of immature mossy cells (Mossy-Calb2) and appearance of mature mossy cells (Mossy-Sv2b cells) was observed; suggesting shared initiating processes.

Thus, our results so far show that the dentate gyrus granule cell lineage proceeds through a sequence of distinct cellular stages, with characteristic and stereotypical spatial distribution from the SGZ to the outer granule cell layer. The early cell types—RGLs, nIPCs and neuroblasts—are clearly distinct states. The presence of a small fraction of dividing RGLs is consistent with self-renewal of the RGL pool, whereas the active cell division of all nIPCs, and their decline over time, support the notion that these cells are a transient amplifying cell type. As granule cells mature, they transition around the third postnatal week from an immature to a mature state, associated with a shift in gene expression involving large numbers of neuronal-related genes. Together, these findings refine and extend previous literature, and we provide extensive lists of cell type-specific gene expression (Fig. 4b, Suppl. Fig. S6).

### Adult neurogenesis resembles early postnatal development

A second fundamental question in postnatal neurogenesis concerns the relationship between embryonic development and adult neurogenesis. To what extent are these processes related? Is adult neurogenesis fundamentally different, relying on a distinct type of adult stem cell which differentiates through distinct cellular states to become mature granule cells, or are they one and the same, with developmental stem cells persisting into adulthood and reiteration of the developmental differentiation program? Notably, since these questions are about similarities, not differences, they cannot be adequately addressed using markers alone (i.e. the fact that a marker is shared does not rule out other molecular differences). In contrast, single-cell RNA-seq strongly limits the possibility of unobserved differences, and thus can reveal similarities with much greater confidence.

In order to address these questions, we generated a third, larger, dataset (Dataset C; 10X Chromium), spanning perinatal (E16.5 – P5), juvenile (P18 – P23) and adult (P120 – P132) animals (Fig. 5a-d, and Suppl. Figs. S1, S7). Notably this dataset extended the range of ages sampled both to embryonic and true adult stages. We again confirmed the previous findings: RGLs, nIPCs and neuroblasts were distinct cell states conserved between juvenile and adult mice, and granule cells matured abruptly around P20. Radial glia-like cells (RGLs) were also confirmed to be clearly distinct from juvenile (and adult) astrocytes (Fig. 5e), expressing particularly transcription factors *Sox4* and *Ascl1*, as well as *Thrsp* (also known as SPOT14), a regulator of adult neurogenesis via lipid metabolism(Knobloch et al. 2013; Knobloch et al. 2014) that is expressed in quiescent radial and non-radial cells that give rise to the neuronal lineage in adult mice.

Overall, the t-SNE visualization revealed a similar process to that of Datasets A and B. We found all the same non-neuronal clusters and GABAergic neurons, and the same trajectory from RGLs to nIPCs, neuroblasts, immature granule cells to mature granule neurons. These clusters also largely expressed the same enriched markers as previously observed (Fig. 5d). However, because the dissection at the earlier timepoints necessarily included some hippocampus proper, we now also observed immature pyramidal neurons (Fig. 5a, grey cells). Interestingly, these cells branched from the granule cell lineage only at the late postmitotic neuroblast stage, indicating a shared differentiation trajectory.

Furthermore, the extension to perinatal timepoints (E16.5, P0 and P5) revealed the existence of a perinatal-specific early neurogenesis programme (Figs. 5b and 5c). Perinatal radial glia (RG) were distinct from juvenile and adult RGLs, with greater expression of *Sox4*, *Sox11* (which are jointly required to generate the hippocampus(Miller et al. 2013)) and *Vim* at the younger ages, and an increased expression of *Notch2* and *Padi2* in the juvenile and adult (Fig 5d). Both cell types were largely quiescent as indicated by the absence of reliable cell cycle genes including *Top2a* and *Cdk1*.

**Figure 5.**
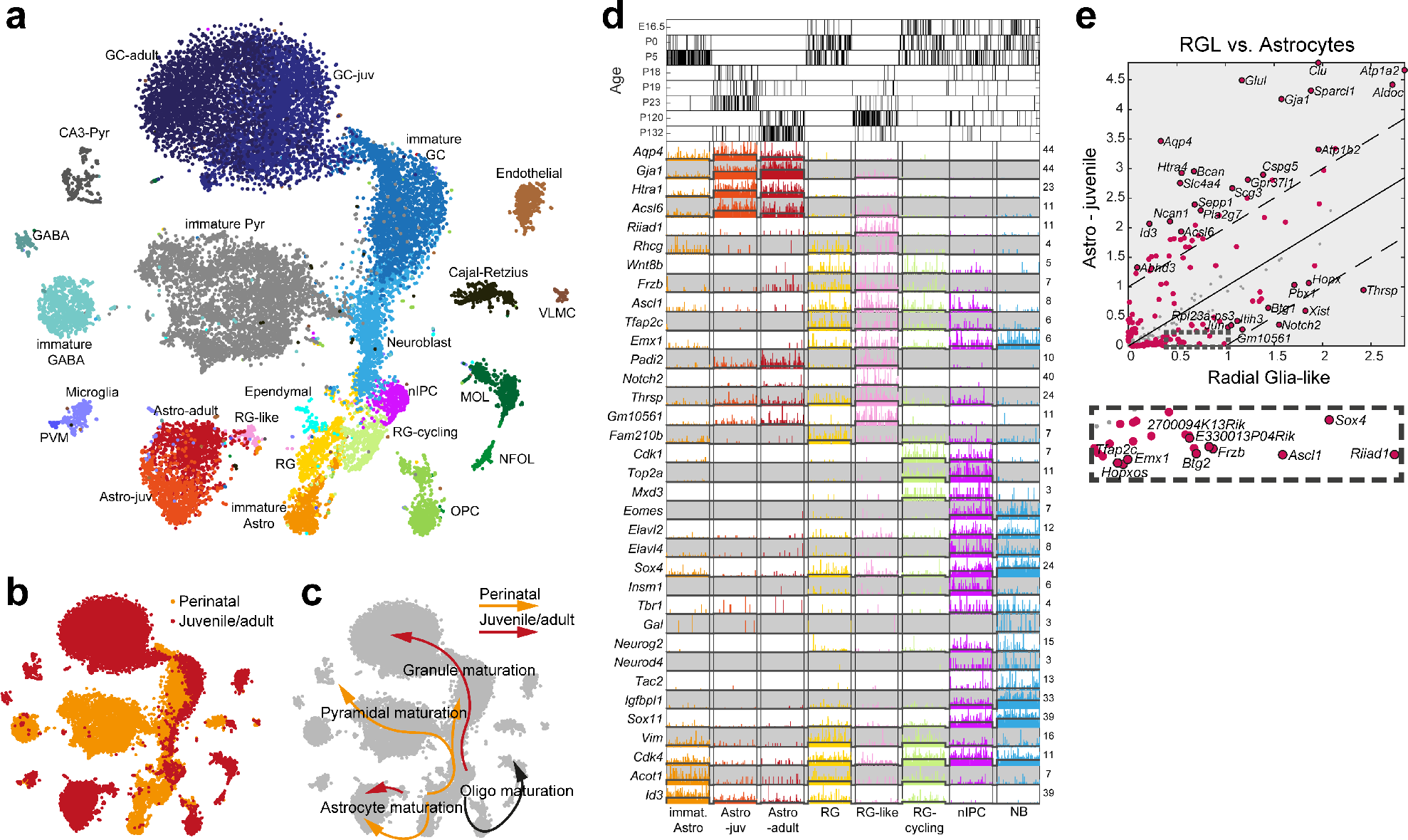
Perinatal, juvenile and adult processes in dentate neurogenesis. (a) tSNE-visualisation of 24,185 cells colored by cluster (24 clusters, names indicated), sampled from perinatal (E16.5-P5), juvenile (P18-23) and adult (P120-132) mice. (b) tSNE colored by sampling age (perinatal – orange, juvenile and adult – red). (c) Developmental trajectories for perinatal (orange) and juvenile/adult (red) maturation processes as suggested in (b) are indicated by arrows. (d) Singe-cell marker gene expression of astrocytes and clusters involved in granule cell neurogenesis. Black bars on top indicate sampling age. (e) Pairwise comparison of gene expression (average number of molecules per cluster) of juvenile/adult RGL versus astrocytes cluster. Red dots are genes enriched at FDR-adjusted q-value < 0.01, with the top-20-significant genes on either side labeled (red dots with stroke). 43 genes were differentially expressed at least two-fold. Solid line at equal expression, dashed lines mark two-fold enrichment.

We also found perinatal-specific cycling radial glia (RG-cycling, Fig. 5d), which differed from nIPCs in their lower expression of neurogenic transcription factors such as *Eomes*, *Elavl2*, *Elavl4*, *Neurog2*, *Neurod4*, *Sox4* and *Sox11*. Although cycling radial glia were only rarely observed after P5, they would be analogous to the cycling radial glia-like nIPCs (Fig. 2e-f) found in juvenile and adult animals, likely representing radial glia-like cells occasionally entering the cell cycle without differentiating. Thus, the stage of initiation of neurogenesis (radial glia and cycling radial glia) showed a clear shift in the second postnatal week from an embryonic form (RG, RG-cycling) to a juvenile and adult form (RGL and cycling radial glia-like nIPCs).

In contrast, nIPCs spanned the full range of ages, from E16.5 to P132 (Fig. 5d, top), as did neuroblasts and immature granule cells. That is, adult forms of these cell types did not generate separate clusters to those of the perinatal or juvenile stages, in contrast to the clear distinction between perinatal and juvenile forms of radial glia. This was true both for cells sampled without selection, and for hGFAP:GFP+ cells sorted by FACS (Supplementary Fig. 1g; compare also the result from Dataset B in Supplementary Fig. 5).

To quantify this observation (Fig. 6), we performed pairwise comparisons of perinatal versus juvenile and adult types (where we would expect significant differences for radial glia only), as well as juvenile to adult (where we would expect minor differences). Comparing perinatal radial glia to juvenile radial glia-like cells (Fig. 6a), we found many genes that were both statistically significantly differentially expressed (red circles) and showed greater than twofold differences (outside the dashed lines), including *Thrsp* in the juvenile and *Vim* in the perinatal. In contrast, comparing juvenile to adult RGLs (Fig. 6d), few genes were significantly differentially expressed and only three (*Pla2g7*, *Sparcl1* and *Aldoc*) showed greater than two-fold change. Thus, radial glia cells, which initiate neurogenesis, switch their molecular identity after P5 and then maintain it until at least P132.

**Figure 6.**
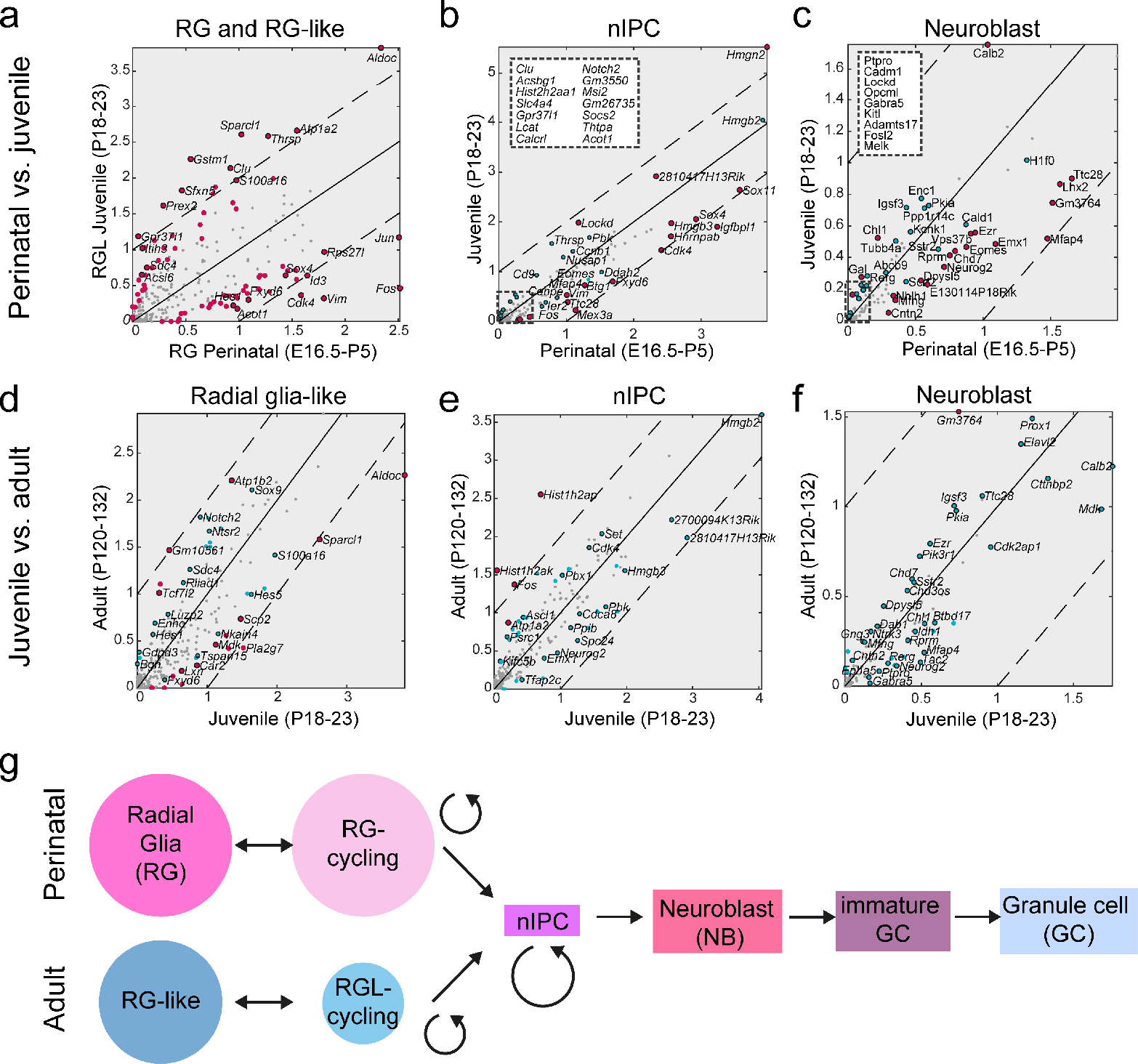
Conservation of differentiation, transformation of initiation. (a-f) Pairwise comparison of gene expression between neurogenesis clusters by sampling age: (a-c) perinatal versus juvenile; (d-f) juvenile versus adult. Red dots are genes enriched at FDR-adjusted q-value < 0.01, with the top-20-significant genes on either side labeled (dots with stroke). Blue dots with stroke are non-significant top-20 genes. Solid line at equal expression, dashed lines mark two-fold enrichment. Number of genes differentially expressed at least two-fold: (a) 16, (b) 2, (c) 1, (d) 4, (e) 3, (f) 0. (g) Suggested model of cell types involved in perinatal and adult dentate neurogenesis, with distinct initiation clusters (radial glia and radial glia-like; size of circle symbolizes frequency), and convergence of the maturation process.

In contrast, both nIPCs and neuroblasts at all ages were near indistinguishable. nIPCs differed by only two genes between perinatal and juvenile (expressed at least two-fold Fig. 6b), and three between juvenile and adult (Fig. 6e). In neuroblasts, not a single neuroblasts gene showed significant two-fold difference between perinatal and juvenile or juvenile and adult mice (Fig. 6c and f). We conclude that although the initiation stage of neurogenesis differs between juvenile and adult compared to perinatal/embryonic, the differentiation trajectory of cells committed to neuronal fate is conserved from the nIPC stage onwards.

One unique feature was observed during development that was absent in the adult. Immature granule cells were found only in the young animals while in the adult these were almost absent (Suppl. Figs. S1, S5d). This was validated *in situ*, where detection of intermediate marker *Fxyd7* was strongly decreased, but not absent, in P24 and P37 (Fig. 4e). This does not necessarily indicate that adult neurogenesis skips a step, but more likely that the great abundance of mature granule cells (which accumulate by age) and the decreasing rate of neurogenesis at this age makes transient intermediate stages more difficult to observe.

## Discussion

We have described the perinatal development and persistent adult neurogenesis of the dentate gyrus using single-cell RNA-seq. Here, we lay out a parsimonious interpretation of dentate neurogenesis supported by our findings.

Our data strongly supports a unified process of early postnatal and adult neurogenesis, with a set of clearly defined, rather than continuous, cell types and transitions. Both the cell types and transitions were invariant over time, but with changes in the number of cells of each type as well as their spatial distribution (Figs. 2 and 5). The long-living stem cell of the dentate gyrus is the radial glia-like cell, as supported by clonal analysis of *Nes*+ RGLs (Bonaguidi et al. 2011). We found that RGLs were predominantly quiescent, even during development (Fig. 2), and in that state they were marked by a very small number of highly enriched markers, which included *Rhcg*, *Vnn1* and *Lpar1*.

After RGLs entered the cell cycle and the nIPC state, a large number of neurogenic transcription factors including *Neurod1*, *Neurod2*, *Neurod4*, and *Eomes* were induced, after which the cells likely would proceed irreversibly to a neuronal fate. However, a small proportion of cycling cells remained RGL-like and did not show robust activation of the neurogenic factors (Suppl. Fig. S4). Thus the pivotal event in dentate neurogenesis is the fate choice occurring as an RGL begins to divide. This fate choice event might involve an asymmetric cell division, or it may be simply stochastic.

Once nIPCs have committed to a neuronal fate, they would either continue to divide, or exit the cell cycle as neuroblasts. The balance between the two would set the amplification gain of the lineage. Data from the SVZ suggests that the average nIPC divides about four times (Ponti et al. 2013), resulting in an approximately eight-fold amplification from RGLs to the neuroblast stage. Since we observed a decrease in actively cycling nIPCs with age – likely explained by a reduced amplification rate – we argue that this also leads to a depletion of neuroblasts. At the same time, this gradual reduction of nIPCs and neuroblasts (Fig. 5), confirms that they must be transient cellular states, while RGLs and astrocytes remained about the same abundance in adult animals. We thus speculate that once neurogenic transcription factors are induced, cells proceed unidirectionally and irreversibly through nIPC, Neuroblast 1 and 2, immature and mature granule cell stages.

Extending the analysis to late embryonic and early postnatal stages (perinatal, E16.5 – P5), we found that the differentiation lineage was unchanged from the nIPC stage through neuroblast and immature granule cell stages, which were indistinguishable between E16.5 and P132. This reinforces the fundamental importance of the induction of neurogenesis, which once induced will slide down a stereotyped and conserved differentiation trajectory. In contrast, on and before P5, including in the embryo at E16.5, the initiation stage of neurogenesis differed markedly compared with the juvenile/adult form. The quiescent radial glia differed, with a perfect separation at the second postnatal week: no perinatal RGs were found after P5, and no juvenile/adult RGLs were found before P18 (Fig. 5d, top). These findings agree with and extend those of (Nicola et al. 2015), who found a shift in the expression of key stem cell marker genes *Sox2*, *Nes*, and *Fabp7* between P7 and P14, indicating a change in the identity of radial glia.

Resequencing at close intervals during postnatal development (Dataset B) revealed another sharp transition during development at P20, when mature granule cells were formed rapidly (a similarly sharp transition was observed for mossy cells). These results indicate that early neurogenesis is reflected by a continuous accumulation of immature granule cells, which at a defined point collectively transit into mature granule cells. Intriguingly, and consistent with the uniqueness of a very special developmental process instructing this transition at around P20, immature granule cells were nearly absent from our sampling after P20 and immature marker *Fxyd7* was detected at drastically lower levels *in situ* from P24. This likely reflects strongly decreased differentiation rates after P20, diminishing the sampling frequency of transient intermediates, as opposed to quiescent progenitor or stable mature populations. A parsimonious interpretation is that before P20, permissiveness for maturation of granule cells is absent, resulting in the accumulation of the intermediate cell type, here defined as immature granule cells.

A shift in the neurogenic profile around P20 is consistent with early studies by Altman and colleagues, where the transition of developmental to adult neurogenesis was seen to occur between P10 and P30 in the rat (Altman & Bayer 1990). In Nestin-eGFP mice, a strong drop in GFP+ neuronal progenitors was reported between P14 and P28 (Gilley et al. 2011). Nicola and colleagues used postnatal time-resolved staining for known markers of neurogenesis and argued that by P14 the adult neurogenic program is essentially in place, since no differences in markers and morphology were observed compared to later time points (Nicola et al. 2015). This partially contradicts our observation of high proportions of neuroblasts and immature types prevailing around P12/P16 with near complete absence of mature cells, compared to after P20. The discrepancy however may lay in the different methods applied. Single-cell RNA-seq is more quantitative than immunohistochemical markers, and resolves gene expression into distinct, data-driven cell types, which is impossible using a few isolated markers. Furthemore, as noted above, markers can detect differences, but not similarities. In order to detect similarities, it is necessary to obtain comprehensive molecular profiles, such as by single-cell RNA-seq.

The sharp transition of immature to mature granule cells following P20 would result in rapid changes of the proportions of cell types, and a transformation of the environment stem cells find themselves in. For example, early postnatal RGLs were surrounded predominantly by other RGLs and neuroblasts, whereas in the adult, nearly all neighbors were mature granule cells, and the environment becomes increasingly less neurogenic compared to early development (Götz et al. 2016). This changing environment is likely the explanation for the reduction in the rate of neurogenesis, leading to the observed reduction of nIPCs and neuroblast numbers. For example, signaling through Notch receptors expressed in neuronal stem and progenitor cells inhibits differentiation to neuronal fates (reviewed in (Louvi & Artavanis-Tsakonas 2006)). We found expression of *Notch1* and *Notch2*, as well as downstream targets *Hes5*, *Sox2* and *Nes* RNA in astrocytes and RGLs, while ligands such as *Jag1* and *Dlk2* were enriched in the mature neuronal states (Suppl. Fig. S6).

In the field of neurodevelopment and adult neurogenesis, major efforts have been invested into isolating and studying populations of stem and progenitor cells, using a wide range of animal models and marker genes (Enikolopov et al. 2015). We showed that several of these genes are not specific to quiescent or cycling progenitors, greatly restricting their utility for such purposes. For instance, *Gfap*, *Hes5* and *Sox2* expression is shared with the closely related Astrocytes, while markers like *Prom1*, *Lpar1* and *Nes* are highly co-expressed by entirely different cell types (endothelial cells and oligodendrocytes) (Fig. 3c), and even *Reln* expression by Cajal-Retzius cells is not exclusive to this population (Suppl. Fig. 2). The wide scope and unbiased nature of our study allowed us to suggest an array of sequential or combinatorial markers that are more narrowly expressed in the relevant cell types and stages of early and late neurogenesis.

One such marker gene is *Tfap2c*, highly specific to RGL and nIPC in our dataset. *Tfap2c* (also known as *Tcfap2c* and *AP2γ*) is a transcription factor that plays a key role in early mammalian extraembryonic development and organogenesis. A murine knockout for this gene is embryonic lethal as early as E7.5. (Lawson & Wilson 2016; Kuckenberg et al. 2012). Recently, *Tfap2c* was shown to be involved directly in hippocampal neurogenesis and suggested to regulate the key neurogenic genes (Schemmer et al. 2013; Pinto et al. 2009; Mateus-Pinheiro et al. 2016). Our data suggest that *Tfap2c* is an exclusive marker for the hippocampus stem-cell niche and we speculate that one role could be to ensure a certain level of proliferation (Kim et al. 2016; Kang et al. 2016) while at the same time determining the fate of subgranular zone RGLs by pre-specification. This would be similar to its function in the developing cortex where it specifies onset of a Tbr2/NeuroD neurogenic lineage program specifically generating certain layer II/III neurons (Pinto et al. 2009).

In summary (Fig. 6g), our findings establish the essential unity of perinatal, juvenile and adult neurogenesis in the mammalian hippocampus, but with sharp changes at around the second (RGLs) and the third (granule cells) postnatal week; demonstrate that neurogenesis proceeds through distinct discrete states; show that the intermediate progenitor and neuroblast stages are molecularly conserved from E16.5 to P132; provide markers with high specificity for each state; show their spatial distribution during development; and suggest a simple model for hippocampal neurogenesis driven by the choice RGLs make to differentiate, or not, as they divide.

## Materials and Methods

### Animals

Male and female wild type CD-1 mice (Charles River) and C57Bl/6 mice, as well as hGFAP-GFP reporter mice (Zhuo et al. 1997) on embryonic day E16.5, as well as between postnatal days P0-132, were used. All experimental procedures followed the guidelines and recommendations of Swedish animal protection legislation and were approved by the local ethical committee for experiments on laboratory animals (Stockholms Norra Djurförsöksetiska nämnd, Sweden).

### Single cell dissociation

Dissection and single cell dissociation were carried out as described before (Marques et al. 2016). Briefly, mice were sacrificed by an overdose of Isoflurane, followed by transcardialperfusion through the left ventricle with artificial cerebrospinal fluid (aCSF, in mM, 87 NaCl, 2.5 KCl, 1.25 NaH_2_PO_4_, 26 NaHCO_3_, 75 sucrose, 20 glucose, CaCl_2_, MgSO_4_). The concentrations of CaCl_2_ and MgSO_4_ were adapted according to sampling age: 2mM and 2mM for <P18, 1mM and 7mM ≥P18. Crucially, aCSF was equilibrated in 95%O_2_ 5%CO_2_ before use and cells were kept on ice or 4°C at all steps except enzymatic digestion. The brain was removed, 300μm vibratome sections were collected, and the dentate gyrus was microdissected. Single-cell suspensions were prepared using the Papain kit (Worthington) with 25-35min enzymatic digestion (depending on age), followed by manual trituration using a BSA-coated p1000 pipette or fire-polished Pasteur pipettes.

### FACS sorting

hGFAP-GFP mice were sacrificed and single cell suspensions prepared as described above. On a BD FACSAria II, GFP-positive cells were sorted into oxygenated aCSF at 4°C, inspected for viability counted, and loaded on the C1 Fluidigm or 10X Chromium, described below.

### Fluidigm C1

For processing on Fluidigm C1, cells were resuspended in aCSF with 5% DNase, counted, and diluted to 500-800 cells/μl for Cell Load at 4°C on a medium-sized chip. All capture sites were imaged and manually checked for the presence of a single, healthy cell; other wells were excluded from library preparation. cDNA synthesis with Unique Molecular Identifiers (UMIs) and 5’ sequencing library preparation by tagmentation followed by multiplexing were carried out using the C1-STRT protocol, as described before(Zeisel et al. 2015). An overview of Fluidigm C1 experiments is found in Supplementary Table 6.

### 10X Genomics Chromium

Dataset A sampling was carried out on two experimental days (Suppl. Table 5), each day pooling samples from male and female mice in a single experiment, to further reduce the risk of batch effects. The dentate gyrus of male (day 1, P35; day 2, P24) and female (day 1, P12; day 2, P16) mice were dissected and dissociated separately. Suspensions were pooled at equal ratios after visual inspection and counting. Dataset C was sampled as indicated in Supplementary Table 5. 300-1000 cells/μl were then resuspended in aCSF and added to the RT mix to aim for sampling of around 3500-5000 cells. All downstream cDNA synthesis (13 PCR cycles), library preparation and sequencing were carried out as instructed by the manufacturer (10X Genomics Chromium Single Cell Kit Version 1 (Dataset A) or Version 2 (Dataset C)).

### Sequencing, QC, data analysis

Libraries were sequenced on the Illumina HiSeq2000, 2500 or 4000 to an average depth of approximately 56,700 (raw, Dataset A), 200,000 and 43,000 (mapped, Dataset B and C) reads per cell, yielding on average (approximate) 2,360 (A), 6,660 (B) and 3,650 (C) distinct molecules, and 1,340 (A), 2,580 (B) and 1,640 (C) genes per cell. Fluidigm C1 library reads were aligned and converted to mRNA molecule counts as described previously (Zeisel et al. 2015). 10X Genomics Chromium samples were aligned to the reference genome and converted to mRNA molecule counts using the “cellranger” pipeline (Dataset A: v 1.1,Dataset C: v 1.3) provided by the manufacturer.

### Data analysis 10X Chromium Dataset A and C, clustering and visualization

10X Chromium Dataset A (Dataset C) datafiles were loaded and merged to one dataset. For more uniform representation of time points, Dataset C datafiles were downsampled as indicated in Suppl. Table 5 (‘Cells_analysed’). Valid cells were defined to have >600 (C: >800) total genes, between 800-20,000 (C: 1000-30,000) molecules, and a ratio of molecules to genes >1.2. This resulted in 6,090 (C: 25546) cells. Next, we removed 46 (C: 732) doublet cells based on co-expression (>1 molecule) of any pair of the following marker genes: *Stmn2* (neurons), *Mog* (oligodendrocytes), *Aldoc* (astrocytes), *C1qc* (microglia), *Cldn5* (endothelial). In Dataset A, we here allocated cells to their sampling age, see separate section below. In Dataset C, we here normalized the number of total molecules per cell to 5000, rounding to integers. We then performed basic feature selection by removing genes with low or wide expression (less than 20 detected molecules (over all cells), or expressed in more than 60% of the cells). Next, we plotted coefficient of variation versus mean (log *CV* vs. log *mean*), fit a linear function and ranked the genes according to distance from the line, to select top 5,000 differentially expressed genes. To exclude an effect of sex or stress genes on clustering, we removed genes previously found to be involved in either (*Ehd2*, *Espl1*, *Jarid1d*, *Pnpla4*, *Rps4y1*, *Xist*, *Tsix*, *Eif2s3y*, *Ddx3y*, *Uty*, *Kdm5d*, *Rpl26*, *Gstp1*, *Rpl35a*, *Erh*, *Slc25a5*, *Pgk1*, *Eno1*, *Tubb2a*, *Emc4*, *Scg5*). Before clustering, the order of cells and genes was permuted and in Dataset A the total number of molecules was normalized to 10,000 per cell. We then calculated PCA projections with 100 components on the log_2_ centered normalized data, and the correlation matrix between cells on the PCA coordinates. On a correlation-based mutual k-nearest-neighbor (KNN) graph (Dataset A *k* = 20; Dataset B *k* = 30), we then calculated the Jaccard distance between cells (fraction of mutual neighbors). Jaccard distances were transformed to edges connecting cells in a graph object (Matlab “graph” function) and the layout was calculated (Matlab “force” option). ‘Cliques’ (cells connected by edges) of less than 10 cells where removed. This Jaccard KNN graph was clustered using MCL (http://micans.org/mcl/), with MCL parameter 1.25 (C: 1.4), minimum cluster size 5 and maximum cluster size 1000 (C: 3000). This resulted in 40 (C: 46) clusters that we plotted on KNN using layout ‘force’. We next calculated a marker table matrix of enriched genes, using top 20 markers power 0, 0.5, 1. Manual inspection of this marker table was necessary to merge mainly granule cell clusters (oversplit by MCL), split the cluster of mixed PVM and VLMC and remove one cluster of suspected doublets, resulting in 22 (C: 25) final clusters. We updated our marker table matrix based on the final clusters, using 20 top markers power 0, 0.5, 1; 675 (C: 756) genes. The same gene selection was used to calculate a dendrogram using linkage (Ward’s, correlation), correlation matrix (each averaged over clusters) and t-SNE projection of all cells with the following parameters: Dataset A: initial PCA 80, perplexity 50, epsilon 100, correlation distance, 1000 iterations; Dataset C (fast implementation BH-tSNE): initial PCA 50, perplexity 100, theta 0.2.

### Data analysis Fluidigm C1 (Dataset B), clustering and visualization

For C1 data analysis we used exactly the same analysis pipeline described above, except for the following modifications: (1) minimum total molecules set to 1500. (2) For doublets removal we used also *C1ql3* as an additional neuronal marker. Further, the threshold for exclusion was set more stringently to >0 detected molecules. (3) For KNN, *k* = 10. (4) Perplexity in t-SNE was set to 60. The number of clusters after MCL was 39; these were manually inspected and partly merged, resulting in 17 final clusters. Here, we started out with 4425 valid cells, removed 719 suspected doublets, as well as cells belonging to small ‘cliques’, resulting in 2706 cells. Clusters suspected to be low quality (lack of specific marker genes) were removed, resulting in 2303 cells in the final dataset.

### Dataset A: Allocation of sampling age to cell

Dataset A sampling was carried out on two experimental days, each day pooling samples from male and female mice in a single experiment, to reduce the risk of batch effects. Thus, the dentate gyrus of male (day 1, P35; day 2, P24) and female (day 1, P12; day 2, P16) mice were dissected and dissociated separately, but pooled at equal ratios to a single Chromium well. To separate male and female cells (and thus timepoints), the sampling time point (P12, P16, P24 or P35) was assigned to each cell according to sample name (10X43_1 or 10X46_1) and expression of male or female-specific genes: *Ddx3y*, *Uty*, *Eif2s3y* for male-, and *Xist* and *Tsix* for female-derived cells. We thus allocated each cell to female (P12, P16) or male (P24, P37) source (i.e. if the sum of female genes was higher than male genes, the cell was defined as female, and vice-versa). For zero expression of either gene sets, the sex was labeled unknown, but was most likely to be male (P24 or P37), since the expression of female genes (mainly *Xist*) was detected in 85% of cells coming from similar samples of known female-only origin (data not shown). In these cases, we labeled the cells P24* or P37*, and expect about 15% of cells classified false-positive.

### Analysis of early neurogenesis and granule cell maturation

To analyze early neurogenesis (Fig. 2) and granule cell maturation (Fig. 4), we generated separate mutual KNN graphs and lists of peak cluster enriched genes. For this, we loaded data only from relevant cell clusters (early neurogenesis – astrocytes, RGL, nIPC, NB1, NB2, GC maturation – NB2, Granule-immature, Granule-mature). Low or widely expressed genes (expressed in less than 20 cells, or more than 60% of all cells) were filtered, the 5000 most differentially expressed genes selected by plotting log *CV* vs. log *mean* (as above). All cells were normalized to 10,000 molecules. PCA projections were calculated with 100 components, and correlation based on the PCA coordinates was calculated. A mutual KNN graph (*k* = 40) was calculated based on correlation as distance, and weights in mKNN graph were replaced by the Jaccard distance (fraction of shared neighbors). Cliques (connected components) of fewer than 10 cells were removed. The graph was calculated (Matlab “force” option), and original clusters were visualized on the new layout. For the analysis of early neurogenesis without nIPC-specific genes, the same pipeline was used, except for the additional removal of genes significantly specific to nIPCs by rank-sum test (FDR 5%) after feature selection.

To identify enriched genes per peak cluster, we loaded data from relevant cell clusters (as above). For each transition identified by the KNN (eg. astrocytes to RGL), we then calculated significantly expressed genes using rank-sum, and fold enrichment as the log mean average in the population divided by the log mean average in all groups. Significantly expressed genes were defined with *q*-value<0.01 in one of the transitions, fold enrichment >1.5, and followed the requirement that fold enrichment in the first or last population be <0.1 (early neurogenesis) or <0.4 (GC maturation).

### Pairwise cluster comparison

Prior to comparison, cells were normalized to 5000 molecules. To remove non-relevant genes potentially adding noise and false positives in later testing, rough feature (gene) selection was performed, testing clusters of interest against all other clusters. Here, only genes passing FDR 20% after t-test on log_2_ *x* + 1-transformed data were selected. Next, the top 3000 differential genes were selected based on log *CV* versus log *mean* (as above). Finally, we performed gene-by-gene rank-sum testing for each pairwise group, recording *q*-value (FDR corrected *p*-value) and log_2_ *x*-fold change. The number of cells included in each cluster by age is listed in Supplementary Table 7.

### RNAscope

CD-1 mice (Charles River) were sacrificed with an overdose Isoflurane and perfused through the left ventricle with PBS. Brains were immediately dissected out, embedded in OCT on dry ice and stored at −80°C. 10μm cryostat sections were collected, covering the anterior-posterior axis of the dentate gyrus. The sections were either fixed in 4% PFA for 5min, rinsed in PBS and stored at −80°C until staining, or immediately frozen at −80°C and fixed after thawing. RNAScope hybridizations were carried out according to the manufacturor's instructions, using the RNAscope Multiplex Fluorescent (Advanced Cell Diagnostics) for fresh frozen sections. Briefly, thawed sections were dehydrated in sequential incubations with ethanol, followed by 30min Protease IV treatment and washing in PBS. Appropriate combinations of hybridization probes (*Ednrb* 473801-C1, *Cdk1* 476081-C2, *Aldoc* 429531-C3, *Ryr2* 479981-C1, *Dcx* 478678-C2, *Plk5* 479971-C3, *Neurod6* 444851-C1, *Sv2b* 479998-C3, *Gad1* 400951-C3, *Ascl1* 313291-C1, *Tfap2c* 488861-C3, *Eomes* 429641-C1, *Igfbpl1* 488851-C2, *Calb2* 313641-C3, *Fxyd7* 431141-C1, *Prox1* 488591-C2, *Ntng1* 488871-C3, *Lhx1* 488581-C1, *Reln* 405981-C2) were incubated for 2h at 40°C, followed by four amplification steps, according to protocol, DAPI counterstaining and mounting with Prolong Gold mounting medium (P36930, Thermo Fisher Scientific). Images were acquired using a Nikon Ti-E with motorized stage.

### Immunohistochemistry

CD-1 mice (Charles River) were sacrificed with an overdose of Isoflurane, perfused through the left ventricle with PBS, followed by 4% PFA. Brains were dissected out, post fixed in 4% PFA 16h, cryoprotected in 10% and 30% sucrose, embedded in OCT, frozen and stored at −80°C. 16μm sections were cut on a cryostat, covering the anterior-posterior axis of the dentate gyrus, and frozen at −80°C until staining. Before staining, thawed sections were treated with 1x Target Retrieval solution (Dako), according to the manufacturer’s instructions. Primary antibodies (goat anti-GFP 1:1000 (Rockland), rabbit anti-GFAP 1:500 (Dako), rabbit anti-GFP 1:1000 (Molecular Probes), goat anti-AldolaseC 1:500 (Santa Cruz Biotech.), goat anti-PDGFRA 1:200 (R&D)) were incubated O/N, 4°C, in PBS with 5% normal goat or donkey serum, 3% BSA, 0.5% Sodium Azide, 0.3% Triton X-100, followed by washing and incubation with appropriate secondary antibodies (donkey anti-goat Alexa 488, donkey antirabbit Alexa 555, donkey anti-rabbit Alexa 488, donkey anti-goat Alexa 555) 1:1000, O/N 4°C in PBS, Hoechst (Molecular Probes) counterstained and mounted with ProLong Gold (Molecular Probes).

## Data Availability

The datasets generated during the current study is available in the GEO repository, accession GSE104323.

## Code Availability

The code used to perform analyses in this paper is available from the authors upon request.

## References

Altman, J. & Bayer, S.A., 1990. Migration and distribution of two populations of hippocampal granule cell precursors during the perinatal and postnatal periods. The Journal of Comparative Neurology, 301(3), pp.365–381. Available at: http://doi.wiley.com/10.1002/cne.903010304.

Altman, J. & Das, G.D., 1965. Autoradiographic and histological evidence of postnatal hippocampal neurogenesis in rats. The Journal of Comparative Neurology, 124(3), pp.319–335. Available at: http://doi.wiley.com/10.1002/cne.901240303 [Accessed November 22, 2016].

Amaral, D.G., Scharfman, H.E. & Lavenex, P., 2007. The dentate gyrus: fundamental neuroanatomical organization (dentate gyrus for dummies). Progress in brain research, 163, pp.3–22. Available at: http://www.ncbi.nlm.nih.gov/pubmed/17765709 [Accessed February 11, 2017].

Bonaguidi, M.A. et al., 2011. In vivo clonal analysis reveals self-renewing and multipotent adult neural stem cell characteristics. Cell, 145(7), pp.1142–55. Available at: http://www.ncbi.nlm.nih.gov/pubmed/21664664 [Accessed February 8, 2017].

Doetsch, F. et al., 1999. Subventricular Zone Astrocytes Are Neural Stem Cells in the Adult Mammalian Brain. Cell, 97(6), pp.703–716. Available at: http://www.sciencedirect.com/science/article/pii/S0092867400807837 [Accessed April 7, 2017].

van Dongen, S.& Abreu-Goodger, C., 2012. Using MCL to Extract Clusters from Networks. In pp. 281–295. Available at: http://link.springer.com/10.1007/978-1-61779-361-5_15 [Accessed February 14, 2017].

Enikolopov, G., Overstreet-Wadiche, L.& Ge, S., 2015. Viral and transgenic reporters and genetic analysis of adult neurogenesis. Cold Spring Harbor perspectives in biology, 7(8), p.a018804. Available at: http://www.ncbi.nlm.nih.gov/pubmed/26238354 [Accessed February 15, 2017].

Eriksson, P.S. et al., 1998. Neurogenesis in the adult human hippocampus. Nature Medicine, 4(11), pp.1313–1317. Available at: http://www.ncbi.nlm.nih.gov/pubmed/9809557 [Accessed April 9, 2017].

Espósito, M.S. et al., 2005. Neuronal Differentiation in the Adult Hippocampus Recapitulates Embryonic Development. Journal of Neuroscience, 25(44).

Gage, F.H., 2000. Mammalian neural stem cells. Science (New York, N.Y.), 287(5457), pp.1433–8. Available at: http://www.ncbi.nlm.nih.gov/pubmed/10688783 [Accessed April 9, 2017].

Gage, F.H., van Praag, H.& Kempermann, G., 1999. Running increases cell proliferation and neurogenesis in the adult mouse dentate gyrus. Nature Neuroscience, 2(3), pp.266–270. Available at: http://www.ncbi.nlm.nih.gov/pubmed/10195220 [Accessed January 13, 2017].

Garcia, A.D.R. et al., 2004. GFAP-expressing progenitors are the principal source of constitutive neurogenesis in adult mouse forebrain. Nature Neuroscience, 7(11), pp.1233–1241. Available at: http://www.ncbi.nlm.nih.gov/pubmed/15494728 [Accessed April 9, 2017].

Gilley, J.A., Yang, C.-P. & Kernie, S.G., 2011. Developmental profiling of postnatal dentate gyrus progenitors provides evidence for dynamic cell-autonomous regulation. Hippocampus, 21(1), pp.33–47. Available at: http://www.ncbi.nlm.nih.gov/pubmed/20014381 [Accessed April 10, 2017].

Gonçalves, J.T. et al., 2016. Adult Neurogenesis in the Hippocampus: From Stem Cells to Behavior. Cell, 167(4), pp.897–914. Available at: http://linkinghub.elsevier.com/retrieve/pii/S0092867416314040 [Accessed November 14, 2016].

Götz, M., Nakafuku, M.& Petrik, D., 2016. Neurogenesis in the Developing and Adult Brain-Similarities and Key Differences. Cold Spring Harbor perspectives in biology, 8(7), p.a018853. Available at: http://www.ncbi.nlm.nih.gov/pubmed/27235475 [Accessed April 7, 2017].

Gould, E.et al., 1999. Hippocampal neurogenesis in adult Old World primates. Proceedings of the National Academy of Sciences of the United States of America, 96(9), pp.5263–7. Available at: http://www.ncbi.nlm.nih.gov/pubmed/10220454 [Accessed April 9, 2017].

Habib, N.et al., 2016. Div-Seq: A single nucleus RNA-Seq method reveals dynamics of rare adult newborn neurons in the CNS. bioRxiv, pp.1–20. Available at: http://www.biorxiv.org/content/early/2016/03/27/045989.abstract.

Hawrylycz, M.J. et al., 2012. An anatomically comprehensive atlas of the adult human brain transcriptome. Nature, 489(7416), pp.391–399. Available at: http://www.nature.com/doifinder/10.1038/nature11405 [Accessed February 13, 2017].

Hsieh, J., 2012. Orchestrating transcriptional control of adult neurogenesis. Genes & development, 26(10), pp.1010–21. Available at: http://www.ncbi.nlm.nih.gov/pubmed/22588716 [Accessed February 7, 2017].

Kang, J.et al., 2016. TFAP2C promotes lung tumorigenesis and aggressiveness through miR-183- and miR-33a-mediated cell cycle regulation. Oncogene. Available at: http://www.nature.com/doifinder/10.1038/onc.2016.328 [Accessed February 10, 2017].

Kaplan, M.S. & Hinds, J.W., 1977. Neurogenesis in the adult rat: electron microscopic analysis of light radioautographs. Science (New York, N.Y.), 197(4308), pp.1092–4. Available at: http://www.ncbi.nlm.nih.gov/pubmed/887941 [Accessed April 9, 2017].

Kempermann, G.et al., 2004. Milestones of neuronal development in the adult hippocampus. Trends in Neurosciences, 27(8), pp.447–452. Available at: http://www.ncbi.nlm.nih.gov/pubmed/15271491 [Accessed April 9, 2017].

Kempermann, G., Kuhn, H.G. & Gage, F.H., 1998. Experience-induced neurogenesis in the senescent dentate gyrus. The Journal of neuroscience: the official journal of the Society for Neuroscience, 18(9), pp.3206–12. Available at: http://www.ncbi.nlm.nih.gov/pubmed/9547229 [Accessed April 9, 2017].

Kempermann, G., Kuhn, H.G. & Gage, F.H., 1997. More hippocampal neurons in adult mice living in an enriched environment. Nature, 386(6624), pp.493–495. Available at: http://www.ncbi.nlm.nih.gov/pubmed/9087407 [Accessed April 9, 2017].

Kim, W. et al., 2016. TFAP2C-mediated upregulation of TGFBR1 promotes lung tumorigenesis and epithelial-mesenchymal transition. Experimental & molecular medicine, 48(11), p.e273. Available at: http://www.ncbi.nlm.nih.gov/pubmed/27885255 [Accessed February 10, 2017].

Knobloch, M.et al., 2013. Metabolic control of adult neural stem cell activity by Fasn-dependent lipogenesis. Nature, 493(7431), pp.226–30. Available at: http://www.ncbi.nlm.nih.gov/pubmed/23201681 [Accessed July 1, 2017].

Knobloch, M.et al., 2014. SPOT14-positive neural stem/progenitor cells in the hippocampus respond dynamically to neurogenic regulators. Stem cell reports, 3(5), pp.735–42. Available at: http://www.ncbi.nlm.nih.gov/pubmed/25418721 [Accessed July 1, 2017].

Kriegstein, A. & Alvarez-Buylla, A., 2009. The Glial Nature of Embryonic and Adult Neural Stem Cells. Annual Review of Neuroscience, 32(1), pp.149–184. Available at: http://www.annualreviews.org/doi/10.1146/annurev.neuro.051508.135600 [Accessed April 7, 2017].

Kuckenberg, P., Kubaczka, C. & Schorle, H., 2012. The role of transcription factor Tcfap2c/TFAP2C in trophectoderm development. Reproductive BioMedicine Online, 25(1), pp.12–20.

Laplagne, D.A. et al., 2006. Functional Convergence of Neurons Generated in the Developing and Adult Hippocampus J. Macklis, ed. PLoS Biology, 4(12), p.e409. Available at: http://www.ncbi.nlm.nih.gov/pubmed/17121455 [Accessed April 7, 2017].

Lawson, K.A. & Wilson, V., 2016. 3 - A Revised Staging of Mouse Development Before Organogenesis. In Kaufman’s Atlas of Mouse Development Supplement. pp. 51–64.

Louvi, A.& Artavanis-Tsakonas, S., 2006. Notch signalling in vertebrate neural development. Nature Reviews Neuroscience, 7(2), pp.93–102. Available at: http://www.nature.com/doifinder/10.1038/nrn1847 [Accessed February 8, 2017].

Maaten, L. van der & Hinton, G., 2008. Visualizing Data using t-SNE. Journal of Machine Learning Research, 9(Nov), pp.2579–2605.

Malatesta, P.& Götz, M., 2013. Radial glia - from boring cables to stem cell stars. Development, 140(3). Available at: http://dev.biologists.org/content/140/3/483.long [Accessed April 7, 2017].

Malatesta, P., Hartfuss, E.& Gotz, M., 2000. Isolation of radial glial cells by fluorescent-activated cell sorting reveals a neuronal lineage. Development, 127(24). Available at: http://dev.biologists.org/content/127/24/5253.long [Accessed April 7, 2017].

Marques, S.et al., 2016. Oligodendrocyte heterogeneity in the mouse juvenile and adult central nervous system. Science, 352(6291).

Mateus-Pinheiro, A.et al., 2016. AP2γ controls adult hippocampal neurogenesis and modulates cognitive, but not anxiety or depressive-like behavior. Molecular Psychiatry. Available at: http://www.nature.com/doifinder/10.1038/mp.2016.169 [Accessed February 8, 2017].

Miller, J.A. et al., 2013. Conserved molecular signatures of neurogenesis in the hippocampal subgranular zone of rodents and primates. Development (Cambridge, England), 140(22), pp.4633–44. Available at: http://www.ncbi.nlm.nih.gov/pubmed/24154525 [Accessed September 25, 2017].

Nicola, Z., Fabel, K.& Kempermann, G., 2015. Development of the adult neurogenic niche in the hippocampus of mice. Frontiers in Neuroanatomy, 9, p.53. Available at: http://www.ncbi.nlm.nih.gov/pubmed/25999820 [Accessed April 10, 2017].

Noctor, S.C., Martínez-Cerdeño, V. & Kriegstein, A.R., 2008. Distinct behaviors of neural stem and progenitor cells underlie cortical neurogenesis. The Journal of comparative neurology, 508(1), pp.28–44. Available at: http://www.ncbi.nlm.nih.gov/pubmed/18288691 [Accessed February 13, 2017].

Overstreet-Wadiche, L.S., Bensen, A.L. & Westbrook, G.L., 2006. Delayed Development of Adult-Generated Granule Cells in Dentate Gyrus. Journal of Neuroscience, 26(8).

Overstreet-Wadiche, L.S. & Westbrook, G.L., 2006. Functional maturation of adult-generated granule cells. Hippocampus, 16(3), pp.208–215.

Pastrana, E., Cheng, L.-C. & Doetsch, F., 2009. Simultaneous prospective purification of adult subventricular zone neural stem cells and their progeny. Proceedings of the National Academy of Sciences of the United States of America, 106(15), pp.6387–92. Available at: http://www.ncbi.nlm.nih.gov/pubmed/19332781 [Accessed February 10, 2017].

Piatti, V.C., Esposito, M.S. & Schinder, A.F., 2006. The timing of neuronal development in adult hippocampal neurogenesis. Neuroscientist., 12(6), pp.463–468.

Pinto, L.et al., 2009. AP2γ regulates basal progenitor fate in a region- and layer-specific manner in the developing cortex. Nature Neuroscience, 12(10), pp.1229–1237. Available at: http://www.nature.com/doifinder/10.1038/nn.2399 [Accessed February 8,2017].

Ponti, G.et al., 2013. Cell cycle and lineage progression of neural progenitors in the ventricular-subventricular zones of adult mice. Proceedings of the National Academy of Sciences of the United States of America, 110(11), pp.E1045–54. Available at: http://www.ncbi.nlm.nih.gov/pubmed/23431204 [Accessed February 7, 2017].

Schemmer, J. et al., 2013. Transcription Factor TFAP2C Regulates Major Programs Required for Murine Fetal Germ Cell Maintenance and Haploinsufficiency Predisposes to Teratomas in Male Mice Schlatt, S. ed. PLoS ONE, 8(8), p.e71113. Available at: http://dx.plos.org/10.1371/journal.pone.0071113 [Accessed February 8, 2017].

Seri, B.et al., 2001. Astrocytes give rise to new neurons in the adult mammalian hippocampus. The Journal of neuroscience: the official journal of the Society for Neuroscience, 21(18), pp.7153–60. Available at: http://www.ncbi.nlm.nih.gov/pubmed/11549726 [Accessed April 9, 2017].

Shin, J. et al., 2015. Single-Cell RNA-Seq with Waterfall Reveals Molecular Cascades underlying Adult Neurogenesis. Cell Stem Cell, 17(3), pp.360–372. Available at: http://www.sciencedirect.com/science/article/pii/S1934590915003124 [Accessed March 31, 2017].

Steiner, B.et al., 2004. Differential regulation of gliogenesis in the context of adult hippocampal neurogenesis in mice. Glia, 46(1), pp.41–52. Available at: http://www.ncbi.nlm.nih.gov/pubmed/14999812 [Accessed April 9, 2017].

Walker, T.L. et al., 2016. Lysophosphatidic Acid Receptor Is a Functional Marker of Adult Hippocampal Precursor Cells. Stem Cell Reports, 6(4), pp.552–565. Available at: http://linkinghub.elsevier.com/retrieve/pii/S2213671116000837 [Accessed June 24, 2016].

Zeisel, A.et al., 2015. Cell types in the mouse cortex and hippocampus revealed by singlecell RNA-seq. Science, 347(6226).

Zhuo, L.et al., 1997. Live Astrocytes Visualized by Green Fluorescent Protein in Transgenic Mice. Developmental Biology, 187(1), pp.36–42.

